# Succinate Dehydrogenase-Deficient Cancer Cells Have Increased Susceptibility to Ym155 Induced DNA Damage

**DOI:** 10.1101/2025.09.29.679343

**Authors:** Qianjin Guo, Sooyeon Lee, Neali Armstrong, Brian Lim, Rebecca C. Schugar, David Tomz, Haixia Xu, Anzel Zhen, Leor Needleman, Electron Kebebew, Justin P. Annes

**Author notes:** Address correspondence to: Justin P. Annes.

## Abstract

The hereditary pheochromocytoma and paraganglioma (hPPGL) syndrome is caused by inherited mutations in Succinate Dehydrogenase genes (SDHx). Affected individuals are predisposed to developing pheochromocytomas (Pheo), paragangliomas (PGL), renal cell carcinoma (RCC) and gastrointestinal stromal tumors (GIST). Notably, tumors with succinate dehydrogenase subunit B (*SDHB*) deficiency demonstrate increased metastatic risk, for which treatments remain palliative. Hence, discovering novel therapeutic avenues to improve the prognosis for *SDHB*-cancer patients is an urgent need. Here we employed human *SDHB*-deficient UOK269 RCC cells (*SDHB*-KO) and isogenic *SDHB*-reconstituted control cells (*SDHB*-WT) to discover SDH-dependent mitochondria-directed cytotoxic agents. Given the reduced ATP-generating capacity of *SDHB*-KO cells, we hypothesized they would be uniquely sensitive to futile cycle induction with mitochondrial ionophores (2,4-Dinitrophenol (2-DNP), BAM15, Niclosamide, Nitazoxanide). Indeed, these compounds exhibited preferential cytotoxicity toward *SDHB*-KO cells. However, the chemotherapeutic compound Ym155 demonstrated the most potent and dramatic (five-fold) preferential cytotoxicity towards *SDHB*-KO cells. Importantly, the SDH-dependent cytotoxicity of Ym155 was validated in both primary human pheochromocytoma cells and mouse pheochromocytoma (MPC) cells. Furthermore, because few SDH-deficient cell lines are available, we buttressed our findings in additional relevant cell lines by modeling SDH-deficiency using chemical SDH enzyme inhibition with 3-nitropropionic acid (3-NPA). We observed a persistent cooperativity between SDH-deficiency and Ym155 cytotoxicity across multiple cell lineages and disease models. Mechanistically, Ym155-induced cytotoxicity was independent of its primary target, Survivin. Instead, SDH-deficiency sensitized cells to Ym155-induced DNA damage. Strikingly, the phenotype of SDH-deficient Ym155 sensitivity was recapitulated by inhibition of the histone demethylase KDM4, a downstream consequence of SDH deficiency. Thus, the accumulation of succinate in SDH-deficient tumors inhibits KDM4 activity, impairs DNA repair and yields enhanced susceptibility to Ym155-induced reactive oxygen species (ROS) generation. The identified intrinsic susceptibilities of *SDHB*-deficient cancers has the potential to be therapeutically leveraged.

## 1. Introduction

Inherited mutations in Succinate Dehydrogenase genes (SDHx) are responsible for the hereditary pheochromocytoma and paraganglioma (hPPGL) syndrome, which includes pheochromocytomas (Pheo), paragangliomas (PGL), renal cell carcinoma (RCC) and gastrointestinal stromal tumors (GIST). Pheochromocytomas and paragangliomas (PPGLs) are neuroendocrine tumors arising from cells of the adrenal medulla and extra-adrenal paraganglia, respectively (*1*). PPGLs are highly hereditary, with over a dozen identified susceptibility genes (*1, 2*). Interestingly, PPGLs can be segregated into three major clusters based upon various molecular characteristics, including transcriptional signature. Cluster 1 tumors are characterized by activation of pseudohypoxia signaling pathways, while clusters 2 and 3 are associated with MAPK and PI3K/mTOR signaling pathways, and the Wnt signaling pathway, respectively (*3, 4*). Cluster 1 is primarily comprised of tumors with germline loss-of-function mutations in succinate dehydrogenase-related genes (SDHx: *SDHA*, *SDHB*, *SDHC*, *SDHD*, and *SDHAF2*), which cause the autosomal dominant hPPGL tumor syndrome (*5*). Inherited pathogenic *SDHB* mutations also confer a 5-14% RCC risk (*6, 7*). Despite distinct cell linages-of-origin, *SDHB*-RCC exhibit similar cardinal molecular features as *SDHB*-PPGLs, including loss of 1p (*8*), succinate accumulation (*9*), genome hypermethylation (*10*) and a pseudohypoxic transcriptional signature (*11*).

Among the SDHx genes, *SDHB* may be considered clinically the most significant given its relatively high prevalence and the aggressive nature of *SDHB*-deficient tumors, with frequent metastases and high mortality (*12–16*). Currently, metastatic SDH-deficient PPGLs are treated with surgical removal of the primary tumor (when possible) and systemic treatment with chemotherapy (e.g., cyclophosphamide, vincristine, dacarbazine), tyrosine kinase inhibitors (sunitinib, Cabozantinib, Axitinib, pazpanib), radiopharmaceutical medications (^131^I-mIBG) or peptide receptor radionuclide therapy (PRRT, ^177^Lu-DOTATTE) (*17*). Unfortunately, these strategies typically demonstrate variable and limited efficacy (*18–20*). Likewise, effective treatment options for metastatic *SDHB*-deficient RCC and GIST, are also limited (*21, 22*). Notably, belzutifan-a selective HIF-2α inhibitor-was recently approved for metastatic PPGL, but at this time published data on efficacy are pending (*23*). Therefore, discovering novel therapeutic avenues that improve the prognosis for metastatic *SDHB*-mutant cancer patients remains an unmet need.

*SDHB* encodes a subunit of the mitochondrial succinate dehydrogenase complex which functions at the intersection of the tricarboxylic acid (TCA) cycle and the electron transport chain (ETC; Complex II). In the TCA cycle, SDH catalyzes the conversion of succinate to fumarate, extracting energy in the form of FADH₂ for the ETC (*24*). In the ETC, the SDHB subunit plays a crucial role as an electron conduit within the SDH complex: its three Fe–S clusters accept electrons from the reduced FADH₂ and pass them to ubiquinone (coenzyme Q) bound to SDHC/SDHD, thereby reducing Q to ubiquinol (*24*). Through this process, SDH links the metabolic breakdown of nutrients to the production of ATP – effectively bridging the TCA cycle with oxidative phosphorylation in the ETC. As a result, SDHB loss disrupts oxidative phosphorylation and leads to accumulation of succinate, which acts as an oncometabolite (*1, 25*). Additionally, loss of *SDHB* and disruption of complex II triggers excess mitochondrial reactive oxygen species (ROS) generation and increases redox imbalance that has been associated with dysregulated iron and copper homeostasis (*26, 27*). Furthermore, accumulated succinate, similar to other intermediate metabolites and byproducts of the TCA cycle, like fumarate and 2-hydroxyglutarate (2-HG), acts as an oncometabolite. The metabolites competitively inhibit the enzymatic activities of α-ketoglutarate–dependent dioxygenases, such as prolyl-hydroxylases (PHD) and JMJD2 family, Ten-eleven translocation (TET) DNA demethylases (*28–31*). As a result, intracellular succinate accumulation causes stabilization of hypoxia-inducible factors (HIFs), epigenetic remodeling, and impaired DNA repair, which collectively promote tumor progression.

Given the impairment of mitochondrial redox homeostasis and ATP generation that occurs as a result of SDH-disruption, therapeutically targeting essential mitochondrial functions might offer therapeutic strategies for treating these cancers. For instance, use of high-dose ascorbic acid to enhance ROS generation has been explored as a potential therapy for *SDHB*-mutant PPGLs (*25, 27, 32*). However, focused research efforts aimed toward exploiting the broader mitochondrial dysfunction of *SDHB*-deficient cells have not been pursued. To address this gap, we investigated whether the mitochondrial dysfunction of *SDHB*-deficient cells can be therapeutically leveraged. By screening a collection of mitochondrial metabolism-targeted small molecules, we identified Ym155 as a highly cytotoxic compound with enhanced activity towards *SDHB*-deficient cells.

## 2. Methods

### 2.1 Cell Culture

The UOK269 cell lines were provided by Dr. W Marston Linehan and maintained in DMEM (Cytiva, SH30243.01) with addition of 10 % fetal bovine serum (FBS,Cytiva,SH30396.03HI), 100 U/mL penicillin-streptomycin (Corning, 30-002-Cl) and 1X Non-Essential Amino Acid (NEAA) Solution (Cytiva,SH30238.01). The MPC cell line was provided by Dr. Arthur S. Tischler and were maintained in DMEM with 10% FBS and 100 U/mL penicillin-streptomycin. Both cell lines were demonstrated to be mycoplasma negative for all experimentation.

### 2.2 Transmission Electron Microscopy

UOK269 (*SDHB*-KO) and UOK269 reconstituted with *SDHB* (*SDHB*-WT) cells were fixed in 2% glutaraldehyde and 4% paraformaldehyde in 0.1 M sodium cacodylate buffer, pH 7.4, at room temperature for 1 hour, followed by post-fixation in 1% osmium tetroxide. Samples were dehydrated in a graded ethanol series (50%,70%,95% and 100%), embedded in EMbed-812 resin, and sectioned using an ultramicrotome. Ultrathin sections (80 nm) were stained with 3.5% uranyl acetate and Sato’s Lead Citrate and imaged using a JEOL JEM-1400 120kV. Mitochondrial area was quantified from randomly selected images using ImageJ software.

### 2.3 Immunofluorescence Staining

Cells were cultured in 96-well plate, treated as indicated, and fixed with 4% paraformaldehyde for 10 minutes. After permeabilization with 0.1% Triton X-100 and blocking with 1% donkey serum, cells were incubated overnight at 4°C with primary antibodies including γ-H2AX (phospho S139) (Abcam Cat # ab11174), H3K9me3 (Active Motif, Cat # 39062), Tyrosine Hydroxylase (Sigma, Cat # AB152) and Phenyl ethanolamine N-methyltransferase (PNMT) (Abcam, Cat # ab119784). After washing, cells were incubated with Alexa Fluor 647–conjugated secondary antibodies for 1 hour at room temperature. Nuclei were counterstained with DAPI. Images were acquired and analyzed on an Operetta CLS High-Content Imaging System using Harmony 4.5 software (Revvity).

### 2.4 MitoTracker Staining

Cells were seeded on 96-well plate and incubated with 100 nM MitoTracker™ Deep Red FM (Invitrogen, Cat # M22425) in complete medium at 37°C for 30 minutes. Following incubation, cells were washed twice with PBS, imaged immediately and analyzed using Operetta CLS High-Content Imaging System using Harmony 4.5 software (Revvity).

### 2.5 Reactive Oxygen Species (ROS) Detection

Mitochondrial ROS were measured using MitoSOX™ Red mitochondrial superoxide indicator (Invitrogen, Cat # M36008). Cells were incubated with 1 μM MitoSOX Red in HBSS at 37°C for 30 minutes. After staining, cells were washed with PBS and analyzed by flow cytometry.

### 2.6 Mitochondrial Membrane Potential (Δψm) Assessment

Δψm was assessed using tetramethylrhodamine ethyl ester (TMRE, Invitrogen, Cat # T669). Cells were stained with 200 nM TMRE in serum-free medium at 37°C for 20 minutes. After washing, cells were imaged and analyzed by Operetta CLS High-Content Imaging System using Harmony 4.5 software(Revvity).

### 2.7 Succinate Quantification by LC–MS

Intracellular succinate levels were measured by liquid chromatography–mass spectrometry (LC– MS). Cells were washed twice with cold PBS and lysed in 80% methanol (pre-chilled to −80°C). After centrifugation at 14,000 × rpm for 10 minutes at 4°C, supernatant was collected for analysis using an Agilent 1290 UHPLC system coupled to an Agilent 6495C triple quadrupole mass spectrometer. Analytes were detected in negative mode using the precursor to product transitions of 117 to 73.1 for succinate and 115 to 71.1 for fumarate. Peak quantification relative to the standard curve was performed using the Agilent MassHunter Qualitative Analysis program.

### 2.8 Western Blotting (WB)

Cells were lysed in RIPA buffer (CST, Cat # 9806S) containing protease and phosphatase inhibitors (Thermo Fisher. Cat # 78440). Protein concentration was determined using a BCA assay. 20 μg protein were separated by SDS-PAGE and transferred to PVDF membranes. Membranes were blocked with 1% BSA in Intercept® Blocking Buffer (LICOR, Cat # 927-60001) and incubated overnight at 4°C with primary antibodies against SDHA (Abcam, Cat # ab14715), SDHB (Abcam, Cat # ab14714), Survivin (CST, Cat # 2808S), γ-H2AX (phospho S139) (Abcam Cat # ab11174), PARP (CST, Cat # 9532T), Caspase-3 (CST, Cat # 14220T), Cadherin-16 (Proteintech Cat # 15107-1-AP) and β-actin (Sigma, Cat # A5316). After incubation with IRDye-conjugated secondary antibodies (1:10000, RT, 1h), blots were imaged with Odyssey system (LICOR).

### 2.9 Cell Viability Assay

Cells were seeded in 96-well plates (10,000 cells per well) and treated with serial dilutions of compounds as indicated. After 72 hours, cells were stained with DAPI and Ethidium Homodimer I (EthD-1, Biotium) for 30 minutes, and imaged using the Operetta CLS High-Content Imaging System using Harmony 4.5 software (Revvity). Viability was determined based on DAPI-positive/EthD-1–negative cells, normalized to vehicle-treated controls. Dose–response curves and IC₅₀ values were calculated using GraphPad Prism software.

### 2.10 Seahorse XF Mitochondrial assay

Mitochondrial respiration was analyzed using the Seahorse XF Analyzer (Agilent) according to the manufacturer’s protocol. Cells were seeded in XF96 plates and incubated overnight. Prior to analysis, cell media was changed to Seahorse XF DMEM supplemented with 1 mM pyruvate, 2 mM glutamine and 10 mM glucose. Basal and maximal oxygen consumption rate (OCR) was measured following sequential injection of oligomycin (1 µM), FCCP (1 µM), and rotenone/antimycin A (0.5 µM each). OCR values were normalized to total protein content per well and analyzed using Wave software.

### 2.11 siRNA-Mediated Knockdown of Survivin

Two different dsRNA were identified in the human *BIRC5* mRNA to selectively target exon 1 (GGACCACCGCAUCUCUACA) or exon 3 (GAGCCAAGAACAAAAUUGC). Cells were transfected with siRNA targeting *BIRC5* or non-targeting control siRNA using Lipofectamine 3000 (Invitrogen, Cat # L3000015) according to the manufacturer’s instructions. After 72 hours, knockdown efficiency was confirmed by Western blot.

### 2.12 CRISPR/Cas9-Mediated KDM4A Knockout

To generate KDM4A knockout cells, a single-guide RNA (sgRNA) targeting human KDM4A was cloned into the pMCB320 lentiviral vector. Lentiviral particles were produced by co-transfecting HEK293T cells with pMCB320-sgKDM4A, pRSV-Rev, pMDLg/pRRE, and pMD2.G plasmids using Lipofectamine 3000. Viral supernatants were harvested 48 hours post-transfection, filtered through a 0.45 μm filter, and used to transduce UOK269 *SDHB*-reconstituted cells (WT) in the presence of 8 μg/ml polybrene. After 48 hours, mCherry-positive cells were sorted by flow cytometry and expanded for downstream validation. KDM4A-gRNA-F 5’-TTGGAGTGAACTGCCTCCAAGAGCGTTTAAGAGC-3’; KDM4A-gRNA-R 5’-TTAGCTCTTAAACGCTCTTGGAGGCAGTTCACTCCAACAAG-3’

### 2.13 Quantitative PCR for mtDNA Copy Number

Total cellular DNA was extracted using the QIAMP DNA blood kit (Qiagen) according to the manufacturer’s instructions. Mitochondrial DNA copy number was assessed by PCR amplification of a conserved mitochondrial genome segment. Quantitative PCR was performed as previously reported(*33*). A total of 15 ng DNA per reaction was used as PCR template. A 221 bp region of human mtDNA was amplified. PCR conditions were as follows: initial denaturation at 94°C for 5 min; followed by 17 cycles of 94°C for 1 min, 58°C for 1 min, and 68°C for 1 min; and a final extension at 68°C for 2 min. PCR products were resolved by gel electrophoresis and quantified for relative mtDNA abundance. Primer sequences were listed as below: Hu_221bp_MitoDNA_F 5’-CCCCACAAACCCCATTACTAAACCCA-3’ and Hu_221bp_MitoDNA_R 5’-TTTCATCATGCGGAGATGTTGGATGG-3’

### 2.14 Bulk mRNA Sequencing (RNAseq)

Total RNA was extracted from UOK269 (KO) and UOK269 reconstituted with SDHB (WT) cells (3 samples each genotype) using the RNeasy Mini Kit. RNA quality was assessed by bioanalyzer. RNA libraries were prepared with an Illumina kit with PolyA selection and sequenced on an Illumina NovaSeq platform to generate 150 bp paired-end reads (PE150), generating ∼30 million paired reads per sample. Raw reads were aligned to the human genome (hg38) using HISAT2, and differential expression analysis was performed with DESeq2. Gene Set Enrichment Analysis (GSEA) was performed in R using the clusterProfiler and enrichplot packages, based on log2 fold-change–ranked gene lists.

### 2.15 Primary human Pheochromocytoma Cell Isolation and Culture

Pheochromocytoma tissue was first placed in a sterile 60 mm Petri dish and immersed in 5 mL of freshly prepared, cold PBS. The tissue was then finely minced into approximately 1 mm³ fragments and transferred to a 50 mL conical tubeand digested by Collagenase-Dispase medium (ThermoFisher, Cat # 17703034). The tube was incubated in a 37°C water bath for 30–45 minutes, with vortexing every 5 minutes to facilitate digestion. If red blood cells are present, the cell pellet is resuspended in 5 mL of red blood cell lysis buffer (Sigma, Cat #, 11814389001). Primary cells were maintained in serum-free DMEM/F-12 (1:1) medium on poly-D-lysine–coated culture dishes to enhance cellular adhesion. To transduce the cells with chromaffin cell marker, cells were transduced with GFP lentiviral construct, driven by human tyrosine hydroxylase (TH) promoter (VectorBuilder), in the presence of 8 µg/mL polybrene to enhance infection efficiency. Following a 24-h exposure, the medium was replaced with fresh DMEM/F-12 (1:1).

### 2.16 Mouse Renal Tubular Epithelial Cell (MRTEC) Model Generation and Culture

Mouse renal tubular epithelial cells (MRTEC) were isolated similarly to prior description (*34*). Briefly, mice were euthanized and perfused with Liver Perfusion Medium (Invitrogen), followed by Liver Digest Medium (Invitrogen). Kidneys were excised, minced into small fragments, and incubated in Liver Digest Medium at 37 °C on a rocker for 20–45 minutes. Digested tissues were gently dissociated by pipetting, washed with Hepatocyte Wash Medium (Invitrogen), and filtered through a 100-μm cell strainer. Cell suspensions were washed again with Hepatocyte Wash Medium, followed by three washes in Dulbecco’s Modified Eagle Medium/Ham’s F-12 (1:1) (DMEM/F-12). The resulting tubular epithelial cell fractions were plated onto collagen I–coated dishes (BD Biosciences) and cultured in DMEM/F-12 supplemented with growth factors (Clonetics, CC-4127). For immortalization, MRTECs were transduced with a doxycycline-inducible SV40 large T antigen (SV40 T-Ag) construct (VectorBuilder), and for drug testing, doxycycline was withdrawn for 7 days prior to Ym155 treatment to ensure the removal of SV40 T-Ag induction.

### 2.17 Statistical Analysis

All quantitative data are presented as mean ± standard error of the mean (SE). No experiment-wide/across-test correction was performed. Statistical comparisons between two groups were performed using unpaired two-tailed Student’s *t*-test unless otherwise noted. For comparisons among multiple groups, one-way or two-way analysis of variance (ANOVA) followed by post hoc test was applied, as appropriate. Dose–response curves and IC₅₀ values were calculated using non-linear regression with a variable slope model (GraphPad Prism v9). A *P* value < 0.05 was considered statistically significant.

## 3. Results

### 3.1 Mitochondrial dysfunction and redox imbalance in *SDHB*-deficient cells

To identify cytotoxic treatments that depend upon SDH-deficiency for efficacy, i.e. SDH-dependent synthetic lethality, we selected human *SDHB*-deficient UOK269 renal cell carcinoma cells (*SDHB*-KO) and isogenic *SDHB*-reconstituted UOK269 cells (*SDHB*-WT) as discovery and control cell-lines, respectively (*35*). Prior to screening, we first characterized the mitochondrial and metabolic phenotype of *SDHB*-KO and *SDHB*-WT cells to ensure utility for our purpose. As anticipated, Western blot confirmed the near absence of SDHB protein in KO cells (minimal non-functional *SDHB* p.R46Q likely present) (Fig. 1A). Consistent with prior work, SDH disruption resulted in significantly increased mitochondrial membrane potential, as assessed by TMRE staining (Fig. 1B and 1C) (*36*). These findings were confirmed with MitoTracker Deep Red FM staining (Supplementary Fig. 1A and 1B), indicating an elevated mitochondrial membrane potential in SDH-deficient cells that reflected metabolic stress. To evaluate mitochondrial oxidative stress, mitochondrial superoxide generation was assessed using MitoSOX fluorescence intensity. Indeed, *SDHB*-KO cells showed higher MitoSOX fluorescence compared to WT cells (Fig. 1D), suggesting higher mitochondrial oxidative stress in KO cells. Next, we evaluated the mitochondrial ultrastructure of KO and WT cells using transmission electron microscopy (TEM). Similar to the human *SDHB*-tumor phenotype, which characteristically exhibits abundant, swollen and clustered mitochondria with degenerate cristae, KO cell mitochondria appeared swollen with increased volume relative to WT cells (Fig. 1E and 1F) (*7, 37*). Collectively, these data indicated that the UOK269 cells faithfully retained key mitochondrial and metabolic phenotypic features of *SDHB*-deficient tumors, which were reversed with forced SDHB expression.

**Fig. 1.**
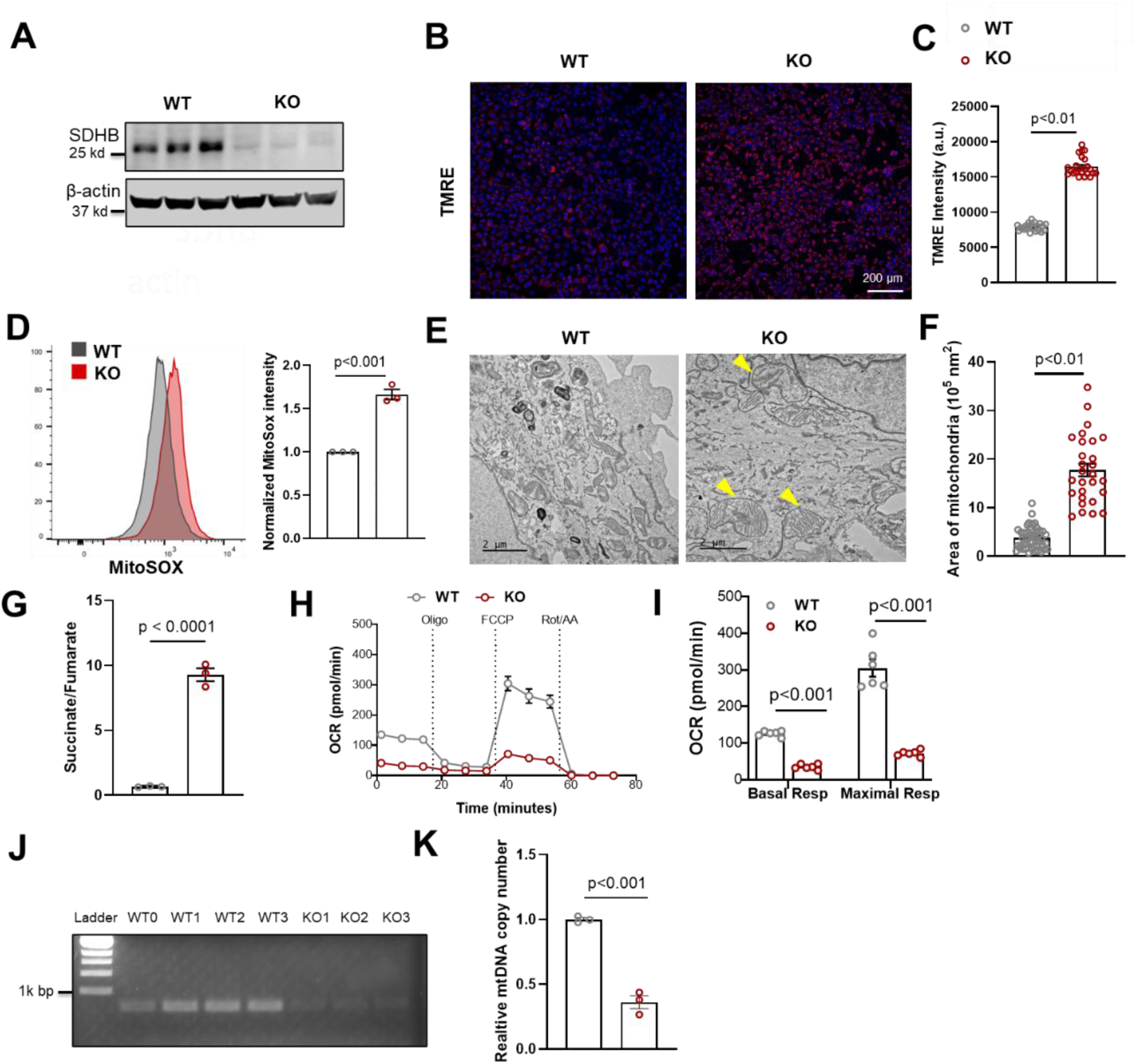
Mitochondrial and metabolic characterization of UOK269 Cells. (A) SDHB expression in *SDHB*-reconstituted UOK269 (WT) and UOK269 (KO) cells was determined by Western blot. β-actin was used as the loading control; n=3 samples. (B) Representative tetramethylrhodamine ethyl ester (TMRE) staining in WT and KO cells. (C) The TMRE average intensity in WT and KO cells determined by Operetta CLS High-Content Analysis System using Harmony 4.5 software (n=3 independent experiments). A linear mixed model analysis was applied in C in which *SHDB* status, and each independent experiment were used as the fixed effect and the random effect, respectively. (D) FACS analysis of MitoSOX-stained WT and KO cells represented as a histogram of florescence intensity (left) and Median Fluorescence Intensity (right; n=3 independent experiments). (E) Transmission electron microscopy (TEM) of WT (left) and KO (right) cells. Characteristic mitochondrial swellings are indicated with yellow arrowheads. (F) Measured mitochondrial area in in TEM images of WT and KO cells (n=47 and 28 mitochondria from >9 representative WT and KO cells). The area was determined and analyzed by ImageJ. (G) Succinate to fumarate ratio was measured in WT and KO cells; n=3 independent experiments. (H) Mitochondrial stress test seahorse profiles of WT and KO cells using an XFe24 Seahorse Analyzer. Oligo, Oligomycin. Rot, Rotenone. AA, antimycin A. (I) Comparison of basal and maximal OCR between WT and KO cells; n=6 independent experiments. (J) Gel electrophoresis analysis of mitochondrial genome fragments generated by PCR, predicting a 221-bp DNA fragment. (K) Measured mitochondrial DNA content from WT and KO cells. WT0, loaded with half the DNA of WT1–WT3, confirmed PCR amplification was within the exponential range for quantification. PCR bands integrated intensity was analyzed by ImageJ; n=3 samples.

Since SDHB is a key component of mitochondrial complex II, we next measured intracellular succinate and fumarate levels by mass spectrometry. KO cells showed a dramatically increased succinate to fumarate ratio compared to WT cells (Fig. 1G). As predicted by the oncometabolite hypothesis, succinate inhibition of alpha-ketoglutarate-dependent enzymes, histone methylation was higher in KO cells (Supplementary Fig. 1C and 1D). Next, to assess *SDHB*-KO cell mitochondrial respiration, oxygen consumption rates (OCR) and complex activities were measured with Seahorse assays. Basal and maximal respiration were reduced in *SDHB*-KO cells and FCCP-treatment failed to induce oxygen consumption (respiratory reserve) (Fig. 1H and 1I); metabolic behaviors characteristic of complex II (SDH) deficiency (*38, 39*). Finally, mitochondrial DNA (mtDNA) copy number was quantified by real-time PCR and was found to be significantly lower in SDHB-KO cells (Fig. 1J and 1K). These findings demonstrate that *SDHB*-deficient UOK269 cells exhibit dramatic *SDHB*-dependent mitochondrial dysfunction and redox imbalance, making them a useful cell system for uncovering SDH-dependent, mitochondrially-targeted synthetic lethality.

### 3.2 Identification of small molecules with preferential cytotoxicity toward *SDHB*-deficient cells

To identify compounds that preferentially target the mitochondrial dysfunction of *SDHB*-deficient cells, we screened a collection of mitochondrial perturbagens for cytotoxicity using *SDHB*-WT and KO cells. Test compounds were strategically curated to target various critical mitochondrial functions (Fig.2A). Overall, we interrogated the hypothesis that SDH-deficiency would increase sensitivity to electron chain disruption or ionophores, excess reactive oxygen species generation or antioxidant depletion, or impaired mitochondrial chaperone function. To test compound cytotoxicity, *SDHB*-KO and -WT cells were seeded into 96-well plates, treated with serial dilutions of each compound (∼[1nM-100uM]), and assessed for viability (72h, DAPI and EthD-1) (Fig. 2B). Cell viability dose–response curves were generated from compound-treated WT and KO cells (Fig. 2C-I; Supplementary Fig. 2B and 2C).

**Fig. 2.**
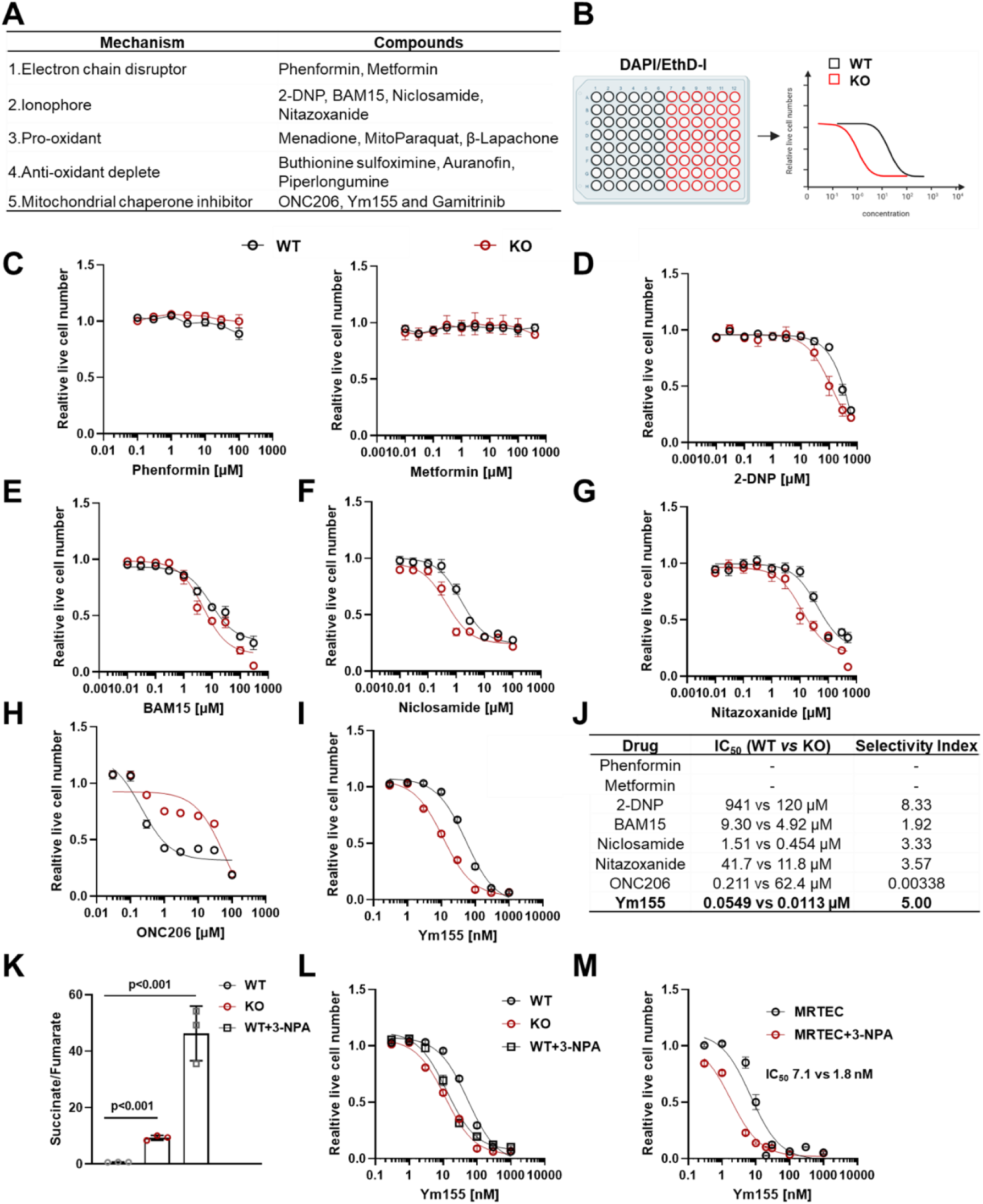
Identification of small molecules selectively targeting *SDHB*-deficient UOK269 cells. (A) Summary of the compounds used in the screen, including known mechanisms of action. (B) Schematic of the experimental workflow. 10,000 UOK269 (KO) and *SDHB*-reconstituted UOK269 (WT) cells were seeded in 96-well plates, treated with compounds for 72 hours, and stained with DAPI and EthD-1 to assess viability. (C–I) Dose–response curves for each compound in WT and KO cells; n=3 independent experiments and at least 3 repeats were performed in each independent experiment. (J) Summary table of IC₅₀ values and selectivity index for compounds tested from (C-I) in WT and KO cells. The selectivity index was calculated as the IC₅₀ ratio of WT to KO cells. (K) Succinate to fumarate ratio in WT, KO and WT cells treated with 100 µM 3-NPA (72h); n=3 independent experiments. Student’s t-test with Bonferroni correction was applied. (L) Dose–response curves of Ym155 in WT, KO, and WT cells treated with 100 µM 3-NPA. n=3 independent experiments. (K) Dose–response curves of Ym155 in mouse renal tubular epithelial cells with or without 100 µM 3-NPA treatment. n=3 independent experiments.

Recently biguanides, including metformin and phenformin, have received attention for their potential to be repurposed as anticancer agents based upon their ability to induce metabolic stress through suppression of mitochondrial ATP generation and induction of ROS generation (*40–43*). However, phenformin and metformin, exhibited minimal cytotoxicity toward either cell type (Fig. 2C). Next, we assessed whether *SDHB*-deficient cells, which have a hyperpolarized membrane potential (Fig. 1C) and reduced ATP-generating capacity, might be more sensitive to futile cycle induction by ionophores (2,4-Dinitrophenol (2-DNP), BAM15, Niclosamide, Nitazoxanide) (Fig. 2D-G) (*44*). Interestingly, these compounds did exhibit the predicted preferential cytotoxicity toward *SDHB* KO cells; however, effects were limited to the [μM] range, potentially limiting *in vivo* application. Next, we interfered with the mitochondrial redox-cycling directly by enhancing ROS generation (Menadione, MitoParaquat, β-Lapachone) or depleting cellular antioxidant capacity (Buthionine sulfoximine, Auranofin, Piperlongumine) (*45–48*). Intriguingly, none of these agents exhibited preferential cytotoxicity toward *SDHB*-KO cells, despite redox imbalance being a typical feature of *SDHB*-KO cells. Next, we tested inhibitors of mitochondrial chaperone functions, including autophagy and apoptosis (ONC206, Ym155 and Gamitrinib) (*49–52*). Unexpectedly, WT cells were more sensitive to ONC206 compared to KO cells while Gamitrinib showed no preferential cytotoxicity (Fig. 2H and Supplementary Fig. 2D). By contrast, among tested compounds, Ym155 exhibited the most potent and selective cytotoxicity toward KO cells, with an effective concentration in the nanomolar range (Fig. 2I and 2J; Supplementary Fig. 2A). Given the potency and selectivity of Ym155, we decided to pursue it as a potential therapeutic agent for *SDHB*-deficient cancers.

To further investigate the SDH-dependence of Ym155 cytotoxicity, we treated *SDHB*-WT cells with a Complex II inhibitor (3-Nitropropionic acid, 3-NPA) to pharmacologically induce SDH deficiency. Importantly, 3-NPA treatment was performed at a concentration ([100μM]) that did not compromise cell viability or cellular replication (Supplementary Fig. 2E). SDH inhibition with 3-NPA treatment was confirmed by induction of the succinate accumulation in WT cells (Fig. 2K). Strikingly, acute SDH-inhibition with 3-NPA-treatment conferred increased SDH-WT cell sensitivity to Ym155, similar to KO cells (Fig. 2L). This observation suggested that Ym155 cytotoxicity was modulated by mitochondrial complex II inhibition and *SDHB* deficiency. To assess whether our findings were specific to tumor-derived UOK269 cells or potentially more general, we established a mouse renal tubular epithelial cell (MRTEC) model from WT mice. Western blotting and RT-PCR confirmed the epithelial identity of the MRTEC cells (Supplementary Fig. 2F and 2G). As anticipated, treatment with 3-NPA resulted in elevated succinate levels in MRTEC cells, reflecting SDH inhibition (Supplementary Fig. 2H). Indeed, pharmacologic inhibition of SDH activity with 3-NPA-treatment increased MRTEC cell sensitivity to Ym155 cytotoxicity (Fig. 2M), further supporting the conclusion that Ym155 cytotoxicity was modulated by SDH activity.

### 3.3 Ym155 cytotoxicity in mouse pheochromocytoma cells (MPC) and primary human pheochromocytoma cells is enhanced by SDH inhibition

Next, we evaluated whether Ym155 exhibited SDH-dependent cytotoxicity towards chromaffin cells, the cellular lineage of most pheochromocytomas and paragangliomas. The mouse pheochromocytoma cell line (MPC) is a well characterized pheochromocytoma cell line (*53–55*), derived from a pheochromocytoma that arose in an NF1 knockout mouse, endogenously expresses phenylethanolamine Nmethyltransferase (PNMT) (*53*). To begin, we confirmed the identity of the MPC cells by demonstrating expression of tyrosine hydroxylase (TH) staining, a marker of catecholaminergic cells (Fig. 3A). Notably a variety of cell-lines used to model pheochromocytomas lack TH expression, a key chromaffin cell identity marker (*56–58*). Having confirmed chromaffin cell lineage, we assessed Ym155 sensitivity of MPC cells with or without SDH inhibition (3-NPA [100 µM]), to evaluate SDH-dependent synthetic lethality. Indeed, SDH inhibition significantly enhanced Ym155-induced cell death, shifting the dose–response curve and lowering the IC₅₀ from 87.22 nM to 11.12 nM (Fig. 3B). These results, consistent with the SDH-dependent cytotoxicity observed in the UOK269 renal cell carcinoma, indicated a persistent cooperativity between SDH-deficiency and Ym155 cytotoxicity across cell lineages.

**Fig. 3.**
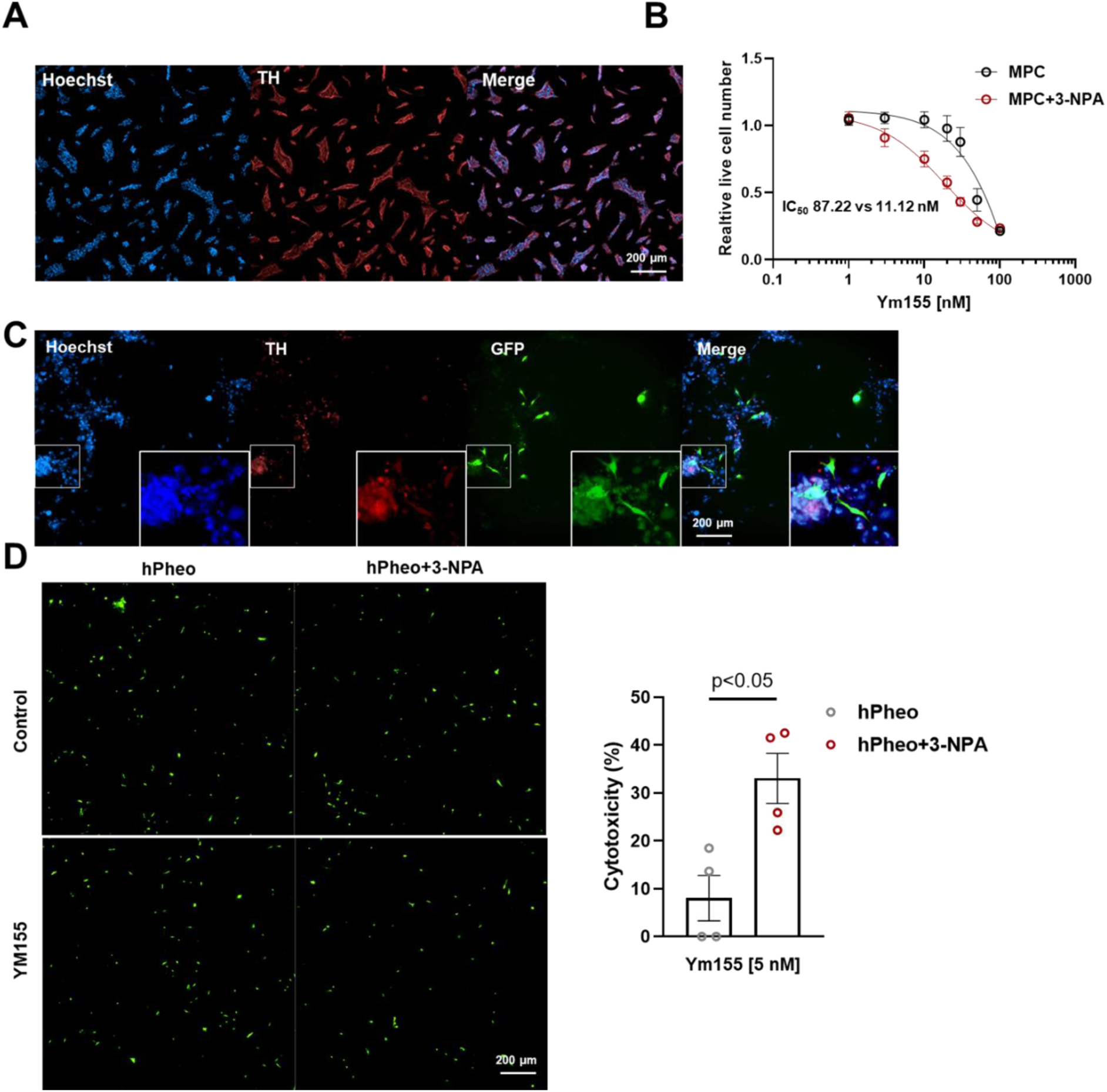
Ym155 cytotoxicity in mouse pheochromocytoma cells (MPC) and primary human pheochromocytoma cells is enhanced by SDH inhibition. (A) Immunofluorescence staining of tyrosine hydroxylase (TH), a marker of catecholaminergic cells, in MPC cells. (B) Dose–response curves of Ym155 in MPC cells treated with or without 100 μM 3-NPA. n=3 independent experiments. (C) Primary human pheochromocytoma cells were transduced with a human TH promoter–driven GFP construct and co-stained with anti-TH antibody. (D) Typical images of GFP positive hPheo cells (left) and quantification analysis of the Ym155 cytotoxicity (right) under 72h treatment in 0 and 5 nM Ym155. 4 repeats were performed.

Next, we extended our findings to a more clinically relevant *in vitro* disease model by assessing Ym155 cytotoxicity toward primary human pheochromocytoma cells. These cultures were established from a 64-year-old woman who developed an apparently sporadic normetanephrine-producing left adrenal gland pheochromocytoma (negative germline SDHx testing and intact SDHB staining on clinical immunohistochemistry). Because primary tumor cultures are compositionally heterogenous, we transduced cells with a human TH promoter-driven GFP construct to identify chromaffin cells and enable targeted analysis. Interestingly, TH-GFP positive cells tended to cluster and form three-dimensional spheroid-like structures, even when plated as individual cells for monolayer culture (black arrowhead, Supplementary Fig. 3A). Co-staining with anti-TH and PNMT antibody confirmed that the TH-GFP reporter reliably identified chromaffin cells (Fig. 3C and Supplementary Fig. 3B). As previously observed, 3-NPA treatment enhanced the Ym155 sensitivity of primary pheochromocytoma cells (Fig. 3D), indicating that compromised mitochondrial SDH function potentiated Ym155 cytotoxicity in primary human pheochromocytoma. However, we note that even in the absence of SDH inhibition, primary human chromaffin cells were highly sensitive to Ym155 treatment (Supplementary Fig. 3C), reducing the observable SDH-dependent effect but, potentially, highlighting a previously unappreciated utility for this late stage therapeutic for pheochromocytoma treatment.

### 3.4 Ym155 induces DNA damage in *SDHB*-deficient cells

Ym155 was originally identified as a suppressor of Survivin (*BIRC5*) expression (*51*). To determine whether Ym155 cytotoxicity toward *SDHB*-deficient cells was mediated *via* Survivin, we examined Survivin protein levels in *SDHB*-WT and *SDHB*-KO cells, with and without exposure to Ym155 (Fig.4A). Western blot analysis revealed that 24-hour Ym155 treatment decreased Survivin levels in *SDHB*-KO cells but not in *SDHB*-WT cells. Interestingly, basal Survivin expression was substantially lower in *SDHB*-KO cells (Fig. 4A), raising a mechanistic question of whether loss of *SDHB* reduces Survivin expression, thereby increasing the vulnerability of *SDHB*-KO cells to Ym155 treatment. To further investigate the regulation of Survivin, we assessed its expression in *SDHB*-WT, *SDHB*-KO, and 3-NPA–treated *SDHB*-WT cells. Indeed, 3-NPA treatment, which inhibits mitochondrial complex II activity, reduced Survivin levels in *SDHB*-WT cells (Fig. 4B), suggesting that SDH dysfunction is associated with decreased Survivin expression and may contribute to the Ym155 vulnerability of *SDHB*-deficient cells.

**Fig. 4.**
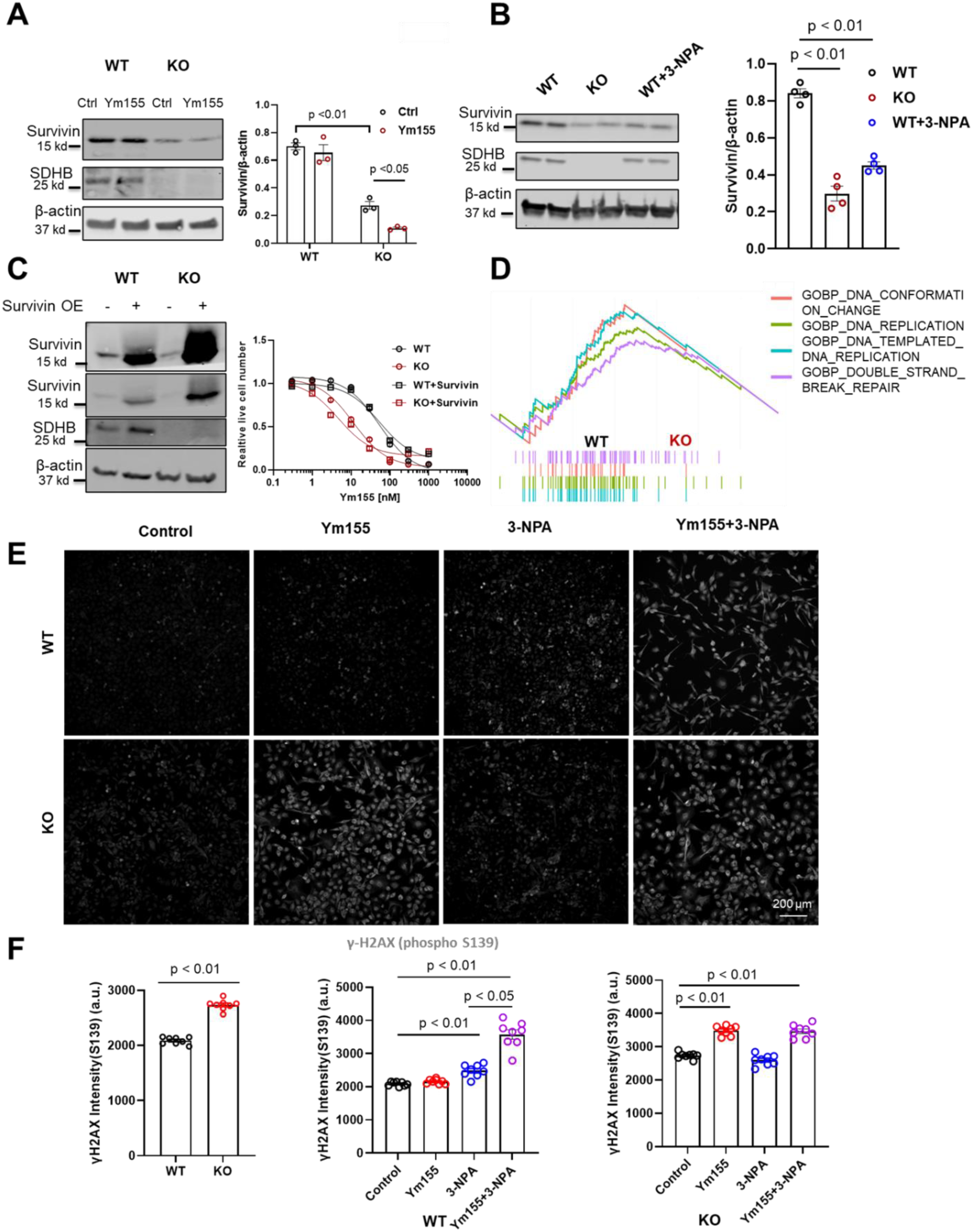
Ym155 promotes cytotoxicity through DNA damage in *SDHB*-deficient cells. (A) Western blot analysis of Survivin expression in UOK269 (KO) and *SDHB*-reconstituted UOK269 (WT) cells following 24-hour Ym155 treatment. n=3 samples. Student’s t-test with Bonferroni correction was applied. (B) Survivin protein levels in WT, KO, and 3-NPA treated WT cells. n=3 samples. Student’s t-test with Bonferroni correction was applied. (C) Typical Western blot image of Survivin overexpression in WT and KO cells (left) and Ym155 dose-curve on WT and KO cells with and without Survivin overexpression (right). n=3 independent experiments. (D) Enrichment plot of selected DSB repair related gene sets in WT and KO cells. (E-F) Immunofluorescence staining of γ-H2AX (phospho S139) and the quantification analysis of γ-H2AX signal in WT and KO cells treated with 10 nM Ym155, 100 µM 3-NPA, or the combination of 10 nM Ym155 and 100 µM 3-NPA for 24 hours. n=3 independent experiments. A linear mixed model analysis was applied in which *SHDB* status and each independent experiment were used as the fixed effect and the random effect, respectively.

To directly test whether Survivin expression controls cellular sensitivity to Ym155 cytotoxicity, we evaluated Ym155 cytotoxicity in *SDHB*-WT and *SDHB*-KO cells which overexpressed Survivin. Notably, overexpression of Survivin did not alter Ym155 sensitivity in either *SDHB*-WT or -KO cells (Fig. 4C). Conversely, RNA interference was applied to knock-down Survivin expression. While knockdown efficiency was confirmed by Western blotting, no significant difference in Ym155 sensitivity was observed between control and Survivin-knockdown cells in either *SDHB*-WT or -KO backgrounds (Supplementary Fig. 4A). Therefore, bidirectional manipulation of Survivin expression did not change the Ym155 sensitivity in either *SDHB*-WT or -KO cells, suggesting that Ym155 cytotoxicity in UOK269 cells was not mediated by Suvivin downregulation. Given that Survivin is an apoptosis-related protein (*59, 60*), we next examined whether Ym155 treatment induced apoptotic signaling. Western blot analysis of PARP/cleaved PARP and Caspase-3/cleaved Caspase-3 after 24 and 48 hours of Ym155 [10 nM] treatment showed no significant changes in either marker (Supplementary Fig. 4A and 4B), indicating that Ym155 cytotoxicity was not associated with PARP/Caspase-3 related apoptosis pathways.

To discover a potential mechanism underlying preferential cytotoxicity of Ym155 towards SDH-deficient cells, we performed bulk RNA sequencing of *SDHB*-WT and -KO cells. Gene set enrichment analysis (GSEA) revealed that double-strand break (DSB) repair gene sets were significantly enriched in *SDHB*-WT cells compared to *SDHB*-KO cells (Fig. 4D and Supplementary Fig. 4D), suggesting that *SDHB*-deficient cells have impaired DNA repair capacity, (*30, 31*). Notably, well-characterized DNA repair related genes (XRCC2, DNA2 and BRIP1) (*61–63*), were downregulated in *SDHB*-KO cells (Supplementary Fig. 4E), consistent with defective DNA double-strand break repair and crosslink repair in these cells. Given these findings, we assessed DNA damage levels by staining for γ-H2AX (phospho-S139), a marker of DSBs (*64*). Indeed, γ-H2AX intensity was increased in *SDHB*-KO cells compared to -WT cells (Fig 4E and 4F). Furthermore, Ym155 treatment significantly increased γ-H2AX signal in *SDHB*-KO cells but not *SDHB*-WT cells. Critically, inhibition of SDH with 3-NPA in *SDHB*-WT cells increased γ-H2AX levels, and the combination of 3-NPA with Ym155 further enhanced γ-H2AX accumulation (Fig. 4E and 4F). These results suggested that SDH dysfunction sensitized cells to Ym155-induced DNA damage.

### 3.5 Pharmacological and Genetic Targeting of KDM4 Enhances Ym155 Sensitivity

To assess the role of mitochondrial dysfunction in Ym155-induced cytotoxicity, we measured mitochondrial ROS levels following Ym155 treatment. MitoSOX staining revealed that Ym155 increased ROS levels in *SDHB*-KO cells but not in *SDHB*-WT cells (Fig. 5A and 5B). Moreover, co-treatment with the non-toxic dose of the ROS scavenger N-acetylcysteine (NAC) significantly rescued *SDHB*-KO cell viability (Fig. 5C and Supplementary Fig. 5A) after 72h Ym155 (10 nM) treatment. These findings indicated that Ym155-induced cytotoxicity was predominantly mediated by mitochondrial oxidative stress in *SDHB*-deficient cells.

**Fig. 5.**
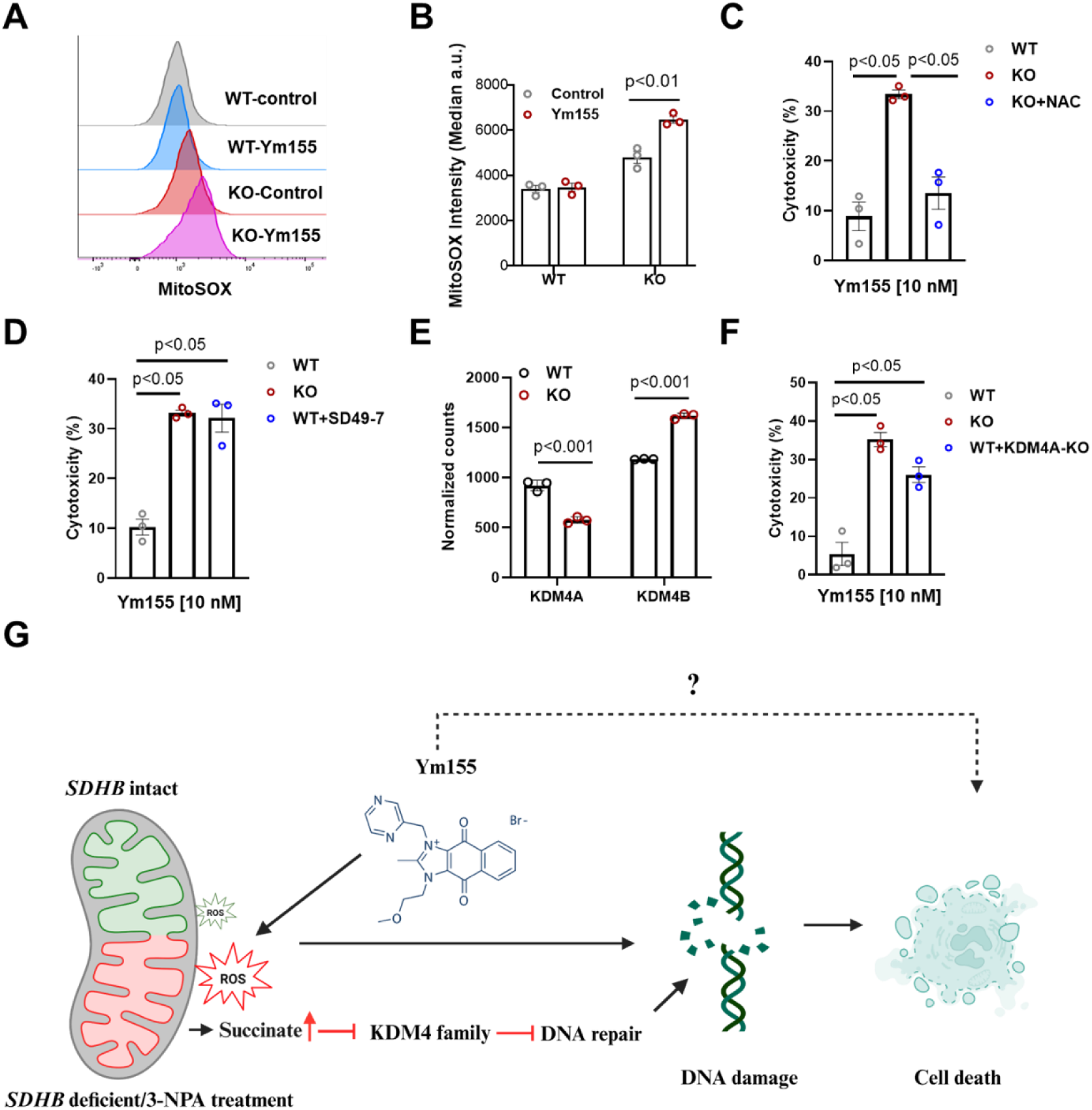
Pharmacological and Genetic Targeting of KDM4 Enhances Ym155 Sensitivity. (A) and (B) FACS analysis showing MitoSOX MFI (Median Fluorescence Intensity) of WT and KO cells with or without 10nM Ym155 24h treatment. n=3 independent experiments. (C) The comparison Ym155 (10 nM,72h) cytotoxicity in WT, KO, and WT cells treated with 3 mM NAC. n=3 independent experiments and each independent experiment include at least 3 repeats. (D) The comparison of Ym155 (10 nM,72h) cytotoxicity in WT, KO, and WT cells treated with 100 nM KDM4 inhibitor SD49-7. n=3 independent experiments and each independent experiment include at least 3 repeats. (E) Transcriptional level of KDM4A and KDM4B in UOK269 and *SDHB*-reconstituted UOK269 cells. n=3 samples. (F) The comparison of Ym155 (10 nM,72h) cytotoxicity in WT, KO and KDM4A knockout cells. n=3 independent experiments and each independent experiment include at least 3 repeats. Student’s t-test with Bonferroni correction was applied in C, D and E.

Succinate accumulation has been reported to impair DSB repair by inhibiting the activity of KDM4 family histone demethylases (*30*). Based on this, we hypothesized that direct inhibition of KDM4 might sensitize *SDHB*-WT cells to Ym155. Treatment of *SDHB*-WT cells with the KDM4 inhibitor SD49-7 at 100 nM, without compromising the cell viability (Supplementary Fig.5B), increased their sensitivity to Ym155 (10 nM) (Fig. 5D), showing a similar cytotoxicity to that of *SDHB*-KO cells. These findings suggested that pharmacologic inhibition of the KDM4 family, enzymes impaired by succinate in the context of SDH-deficiency, mimics the DNA repair vulnerability of *SDHB*-deficient cells.

Since KDM4 inhibition leads to accumulation of H3K9me3, we next evaluated H3K9me3 levels following treatment with 3-NPA or SD49-7. Immunofluorescence staining analysis revealed that both treatments led to an increase in H3K9me3 levels in WT cells (Supplementary Fig.5C), consistent with the predicted inhibition of KDM4 demethylase activity. To identify which KDM4 member is involved, we analyzed the Cancer Genome Atlas (TCGA) datasets for renal cell carcinoma (RCC) and pheochromocytoma/paraganglioma (PPGL) (*11*). KDM4A, but not KDM4B, showed a consistent decrease in *SDHB* tumor samples compared to other mutant tumors (Supplementary Fig. 5D). Consistent with this, we observed a reduction in KDM4A expression in *SDHB*-KO cells, whereas KDM4B expression was increased (Fig. 5E). To determine whether KDM4A plays a functional role in regulating Ym155 sensitivity, we depleted KDM4A in *SDHB*-WT cells (Supplementary Fig.5E). KDM4A loss did not affect either *SDHB*-WT or -KO cell growth (Supplementary Fig.5F), but loss of KDM4A sensitized *SDHB*-WT cells to Ym155 (10 nM) treatment (Fig. 5F). These results indicate that loss of KDM4A increases sensitivity to Ym155 in the context of SDH dysfunction.

## 4. Discussion

In the present study, we identified a conceptually promising therapeutic strategy for *SDHB*-mutated cancers based upon a focused interrogation of mitochondria-targeted small-molecules. Among tested drugs, Ym155 exhibited the highest potency and preferential cytotoxicity toward multiple *SDHB*-deficient (genetic) and SDH-inhibited (chemical) cell models; including primary human pheochromocytoma cells. Historically, drug targets identified *via* synthetic lethality are effective across syndromic cancer types, e.g. PARP-inhibition for BRCA1/2 cancer (breast and ovarian) and HIF2α-inhibition for VHL cancers (RCC, hemangioblastoma, neuroendocrine tumor and PPGL) (*65–67*); broadening the potential impact of this work and raising the possibility for application towards other oncometabolite-driven cancers (e.g. IDH1/2 and FH cancers) (*30*). Such an approach not only highlights a therapeutic strategy tailored to *SDHB*-deficient tumors, but also establishes a paradigm for exploiting vulnerabilities across a wider spectrum of malignancies. Mechanistically, we found that the cytotoxicity of Ym155 was independent of Survivin, a primary reported target (*51*). Instead, cytotoxicity correlated with DNA damage, particularly under conditions of SDH dysfunction, and was modulated by ROS inhibition.

Our findings align with and extend prior studies that have highlighted therapeutic vulnerabilities associated with defective mitochondrial function and impaired DNA repair in *SDHB*-deficient tumors. A finding from our compound screen was that mitochondrial ionophores exhibited preferential cytotoxicity toward *SDHB*-deficient cells. Although we did not pursue this avenue in detail within the scope of the current study, the finding is interesting. Ionophores perturb mitochondrial membrane potential and ion homeostasis, which are already destabilized by SDH loss. Their selective activity therefore raises the possibility that further interrogation of mitochondrial ion flux and bioenergetic stress could uncover novel vulnerabilities in *SDHB*-mutant tumors. Ascorbic acid was shown to induce more oxidative DNA damage in *SDHB* knockdown MPC model (*32*). Additionally, Olaparib, a PARP inhibitor, was found to enhance the cytotoxicity of temozolomide in mice bearing *SDHB* knockdown PPGL allografts, acting through the PARP1-mediated base excision repair (BER) DNA repair pathway (*68*). However, blocking DNA repair using Olaparib showed no viability difference between UOK269 cells (KO) and *SDHB*-reconstituted (WT) UOK269 cells (Supplementary Fig. 5G). These data indicate that UOK269 cells may not rely heavily on homologous recombination repair under basal conditions, but become vulnerable when exposed to exogenous DNA-damaging stress. The lack of effect from PARP inhibition alone, but strong sensitivity to DNA damage, points to a potential threshold effect— where mitochondrial dysfunction predisposes cells to toxic stress only when damage exceeds a certain level.

Ym155 is a compound originally characterized as a transcriptional inhibitor of the apoptosis regulator Survivin and shows broad activity across many tumor types (*69*). However, the mechanism of the Ym155 cytotoxicity is controversial. Increasing evidence suggests that Ym155 induces cytotoxicity primarily through the induction of DNA damage, rather than through suppressing the promoter activity of Survivin and decreasing its expression (*70–72*). Additionally, the AMPK-mTOR axis was also reported as a target of Ym155 in cancer cells (*73, 74*). In our study, Ym155 induced oxidative stress and DNA damage, leading to preferential cytotoxicity in *SDHB*-deficient cells. These effects appear to be particularly potent in the context of SDH loss, where mitochondrial dysfunction and redox imbalance sensitize cells to ROS-mediated DNA damage.

We provided evidence that the DNA damage response in *SDHB*-deficient cells is mediated, at least in part, by the histone demethylase KDM4A, a known α-ketoglutarate–dependent enzyme involved in DNA double-strand break (DSB) repair (*31, 75, 76*). We showed that pharmaceutical inhibition or genetic depletion of KDM4A sensitized *SDHB* WT cells to Ym155, mimicking the phenotype of *SDHB*-deficient cells. Moreover, both 3-NPA treatment and KDM4A inhibition led to elevated H3K9me3 levels, consistent with blocked demethylase activity with succinate accumulation (*28, 30*). These findings suggest that the epigenetic consequences of SDH loss— particularly *via* inhibition of KDM4A—contribute to impaired DNA repair and enhanced drug sensitivity.

Similar to *SDHB*, fumarate hydratase (*FH*) and isocitrate dehydrogenase (*IDH*) mutations also lead to the accumulation of oncometabolites, which promote tumor growth. *FH* mutations cause fumarate accumulation, while *IDH* mutations lead to the production of 2-hydroxyglutarate (2-HG). Interestingly, prior high throughput drug and genetic screens performed with *FH* and *IDH* mutant cell lines to identify the synthetical lethality focused on targets related to cancer metabolism (*77–81*); consistent with the dependence of these metabolically-altered cell mutant cancer cell lines on compensatory pathways and/or specific nutrients. As shown here, the fixed metabolic defect of SDH-deficient cells offers exploitable therapeutic vulnerabilities. In the future, it is promising to perform high throughput drug or genetics screen to identify novel synthetical lethality in *SDHB*-deficient tumor cells.

While our findings uncover a novel therapeutic strategy associated with SDH dysfunction, several limitations should be noted. First, although Ym155 showed robust and reproducible activity in multiple cell-based models, its precise molecular target in the context of mitochondrial dysfunction remains to be fully elucidated. Additionally, although our data indicate that Ym155-induced DNA damage is enhanced in *SDHB*-deficient cells or by KDM4A inhibition, it remains possible that additional stress pathways or off-target mechanisms contribute to this effect (Fig. 5G). Second, most of our mechanistic insights were derived from experiments performed with UOK269 cells. Hence, further validation in patient-derived models and/or *in vivo* systems will be critical to assess translational applicability. Third, Ym155 previously demonstrated limited efficacy in clinical trials for other cancer types, potentially due to poor pharmacokinetics or, perhaps, suboptimal patient / cancer selection (*82–85*). Thus, there may be challenges to evaluating this compound’s utility for treatment of metastatic pheochromocytoma and paraganglioma.

Despite these limitations, our study highlights a promising therapeutic strategy for targeting the metabolic-epigenetic vulnerabilities of *SDHB*-deficient tumor cells. By linking mitochondrial complex II dysfunction to impaired DNA repair *via* KDM4A inhibition and enhanced sensitivity to DNA-damaging agents, we identify a mechanistic basis for therapeutic intervention in a disease with few effective options.

## CRediT authorship contribution statement

**Qianjin Guo:** Writing – review & editing, Writing – original draft, Visualization, Validation, Project administration, Methodology, Investigation, Formal analysis, Data curation, Conceptualization. **Sooyeon Lee:** Writing – review & editing, Visualization, Validation, Formal analysis, Data curation. **Neali Armstrong:** Writing – review & editing, Resources. **Brian Lim:** Methodology, Resources. **Rebecca C. Schugar:** Writing – review & editing, Formal analysis, Resources, Methodology. **David Tomz:** Writing – review & editing, Formal analysis, Resources. **Haixia Xu:** Writing – review & editing, Resources. **Anzel Zhen:** Resources. **Leor Needleman:** Writing – review & editing. **Electron Kebebew:** Writing – review & editing, Supervision, Resources, Funding acquisition. **Justin P. Annes:** Writing – review & editing, Supervision, Project administration, Funding acquisition, Conceptualization.

## Ethical approval

This study was reviewed and approved by the Ethics Committee of Stanford University. All animal experiments were performed in compliance with the Institutional Animal Care and Use Committee and the Stanford University Administrative Panel on Laboratory Animal Care (APLAC).

## Acknowledgements

This work was supported in part by a philanthropic gift from the Lau Family to advance research on SDHB-related disease, and by research grants from the SDHB PheoPara Coalition (JPA) and the Harry A. Oberhelman, Jr. and Mark L. Welton endowment (EK). Additional grant support was received from the NIH NIDDK (JPA: U01DK136965, R01DK101530 and R01DK119955; JPA and LN: T32DK007217), the Neuroendocrine Tumor Research Foundation (QG). The data in Supplementary Figure 4H is part based upon data generated by the TCGA Research Network: https://www.cancer.gov/tcga. Figure 5G was created with BioRender.com

