## Supplementary material for "Succinate Dehydrogenase-Deficient Cancer Cells Have Increased Susceptibility to Ym155 Induced DNA Damage": Supplementary material.pdf

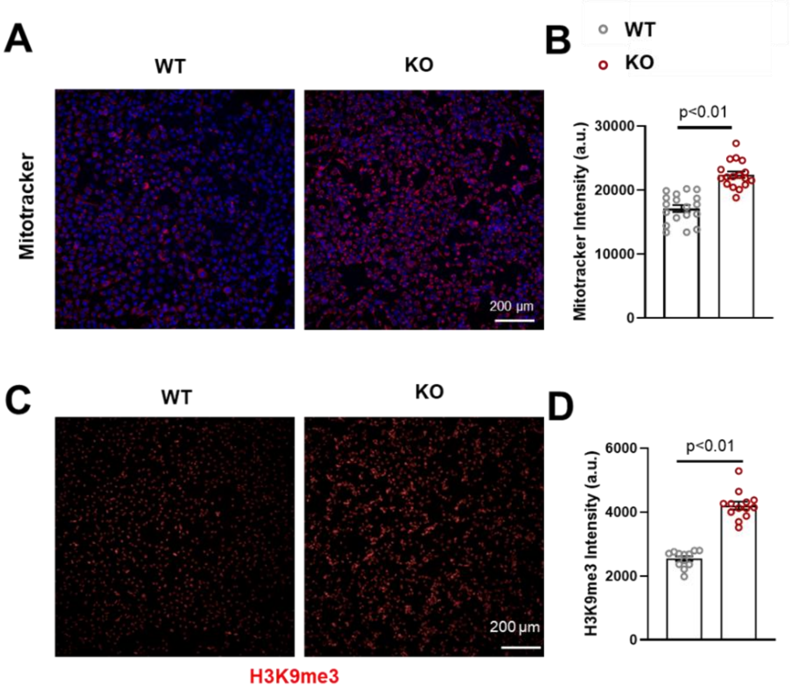


**Supplementary Fig. 1 Characterization of UOK269 (KO) and *SDHB*-reconstituted UOK269 (WT) cells.** (A) Representative Mitotracker staining in UOK269 (KO) and *SDHB*-reconstituted UOK269 (WT) cells. (B) Quantification analysis of Mitotracker average intensity in WT and KO cells, and the intensity analysis was analyzed by ImageJ. n=3 independent experiments. (C) Representative H3K9me3 staining in WT and KO cells. (D) Quantification analysis of H3K9me3 average intensity in WT and KO cells, and the intensity analysis was analyzed by ImageJ. n=3 independent experiments. A linear mixed model analysis was applied in B and D in which *SHDB* status and each independent experiment were used as the fixed effect and the random effect, respectively

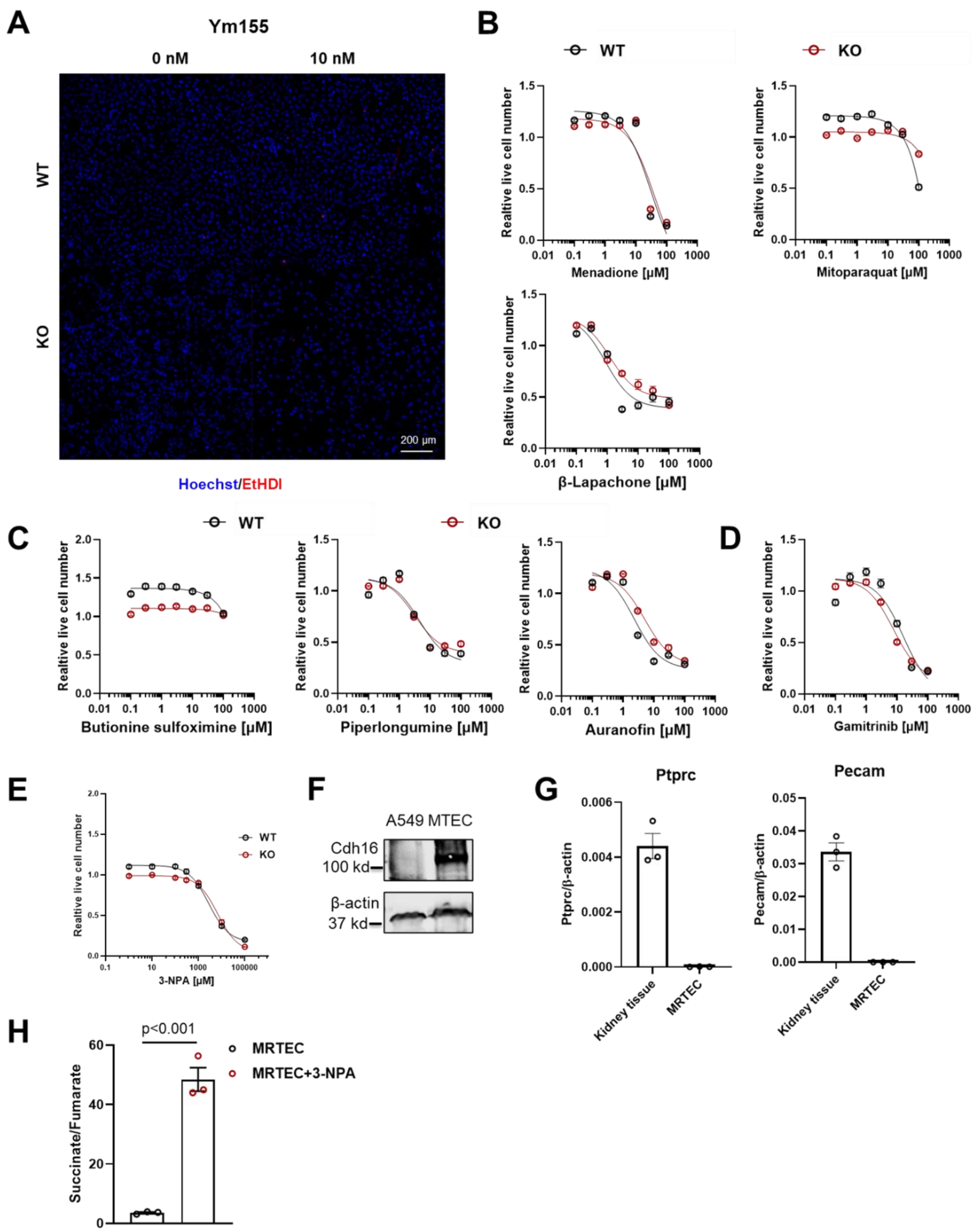


**Supplementary Fig. 2 Compounds screen on UOK269 cells and characterization of MTEC model.** (A) Typical Hoechst and EtHDI staining of UOK269 (KO) and *SDHB*-reconstituted UOK269 (WT) cells under 72h treatment in 0 and 10 nM Ym155. (B-D) Dose–response curves for pro-oxidant, antioxidant depleters and Gamitrinib in WT and KO cells. n=2 independent experiments and 6 repeats were performed in each independent experiment. (E) Dose–response curves for 3-NPA in WT and KO cells. n=3 independent experiments and at least 2 repeats were performed in each independent experiment. (F) Western blot analysis of Cdh16 expression in MRTEC cells and A549 cells. β-actin was used as loading control. (G) RT-PCR analysis of immune cell and endothelial cell marker genes in kidney tissue and MRTEC cells. n=3 samples. (H) Succinate to fumarate ratio in MRTEC cells with and without 100 μM 3-NPA treatment. n=3 independent experiments.

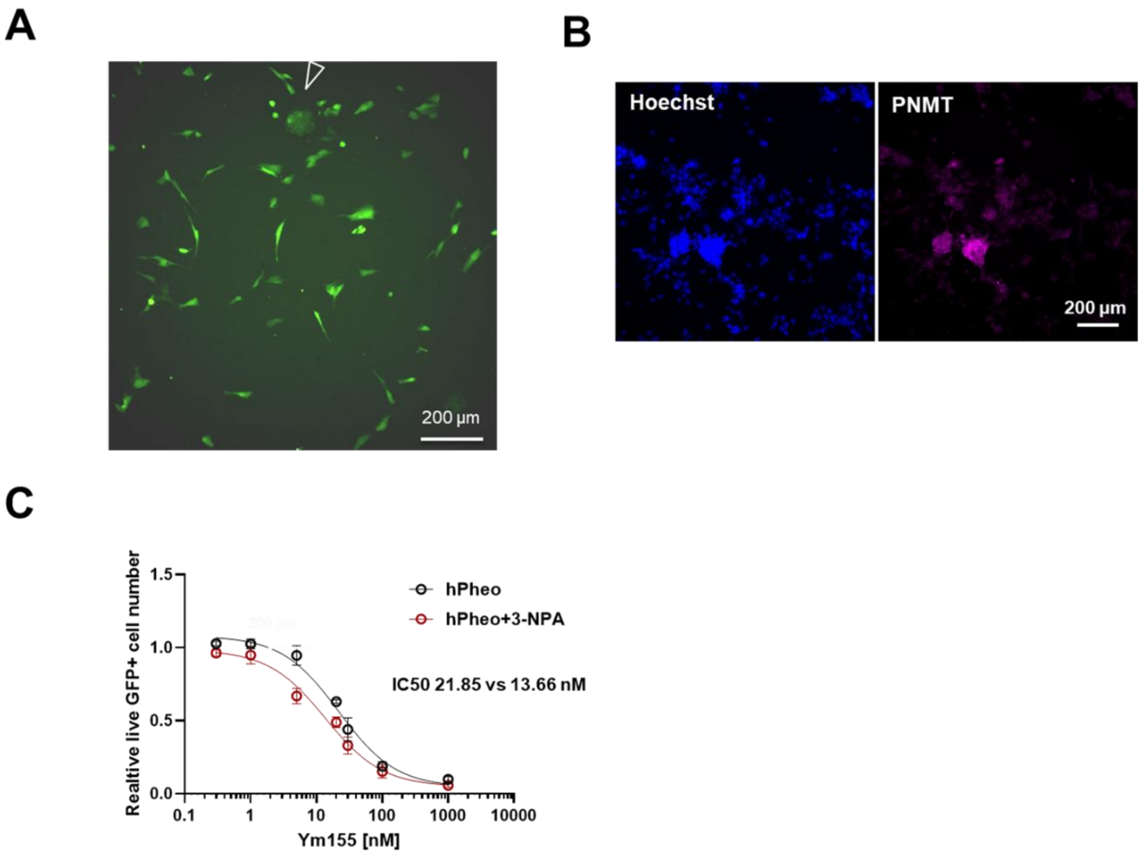


**Supplementary Fig. 3. Characterization of primary human pheochromocytoma cells.** (A) Live-cell imaging by Operetta CLS High-Content Imaging System showed that TH-GFP positive cells form spheroid-like structures. Black arrowhead indicates a typical spheroid-like structure. (B) Immunofluorescence staining of phenylethanolamine N-methyltransferase (PNMT) in primary human pheochromocytoma cells. (C) Dose–response curves of Ym155 in primary pheochromocytoma cells with or without 100 µM 3-NPA treatment. At least 4 repeats were performed at each concentration.

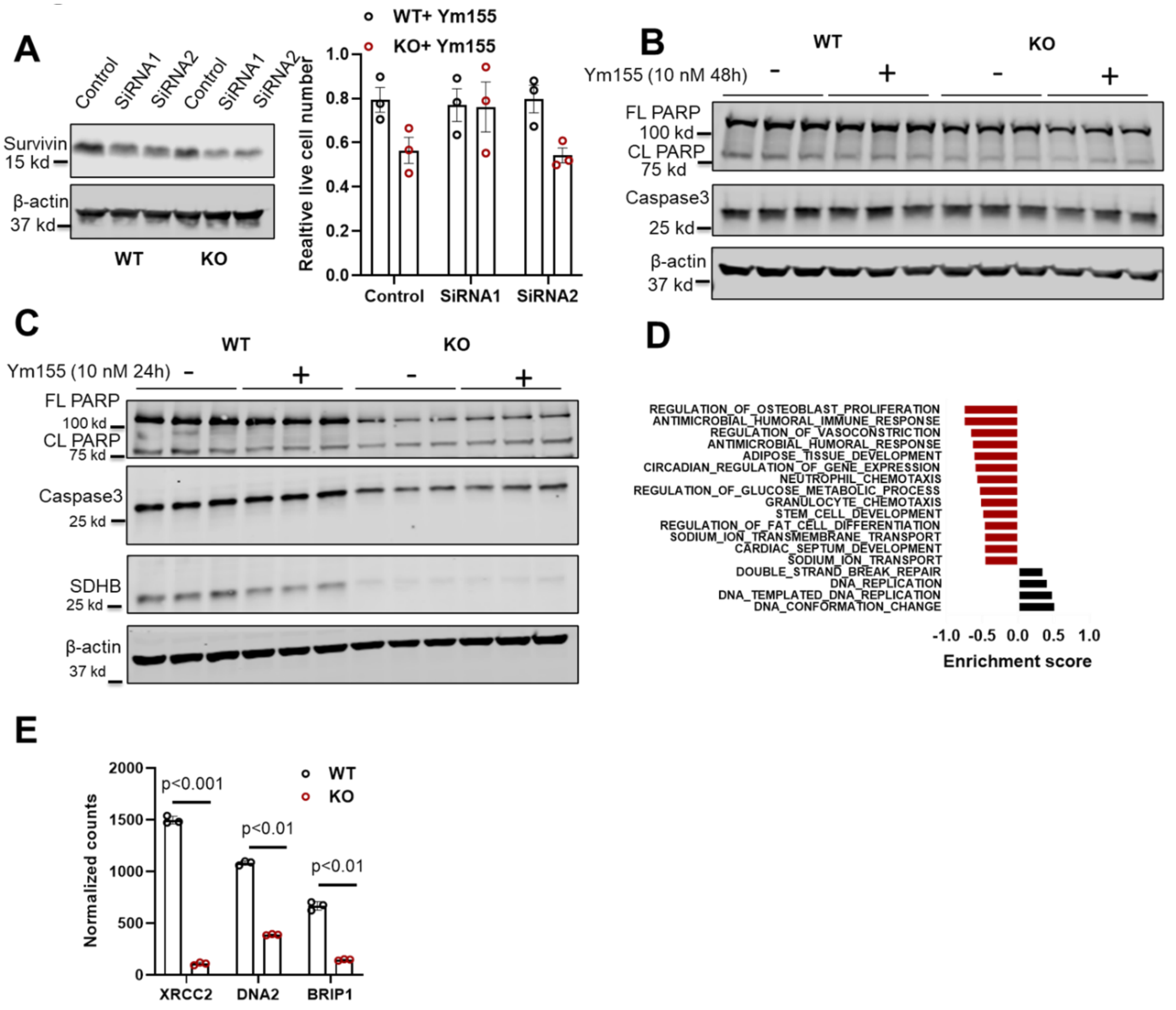


**Supplementary Fig. 4. Ym155 induces DNA damage in a Survivin independent way in UOK269 Cells.** (A) SiRNA medidated Survivin knock down (left) and effect of Survivin knockdown on Ym155 (10 nM, 72h) sensitivity in UOK269 (KO) and *SDHB*-reconstituted UOK269 (WT) cells. n=3 independent experiments. (B-C) Western blot analysis of cleaved PARP and cleaved Caspase-3 WT and KO cells following 24 and 48 hours of Ym155 treatment. n=3 samples. (D) Gene set enrichment analysis (GSEA) showing enrichment of DNA double-strand break (DSB) repair pathways in WT versus KO cells. (E) Transcriptional level of XRCC2,DNA2 and BRIP1 in *SDHB*-WT and -KO cells. n=3 samples.

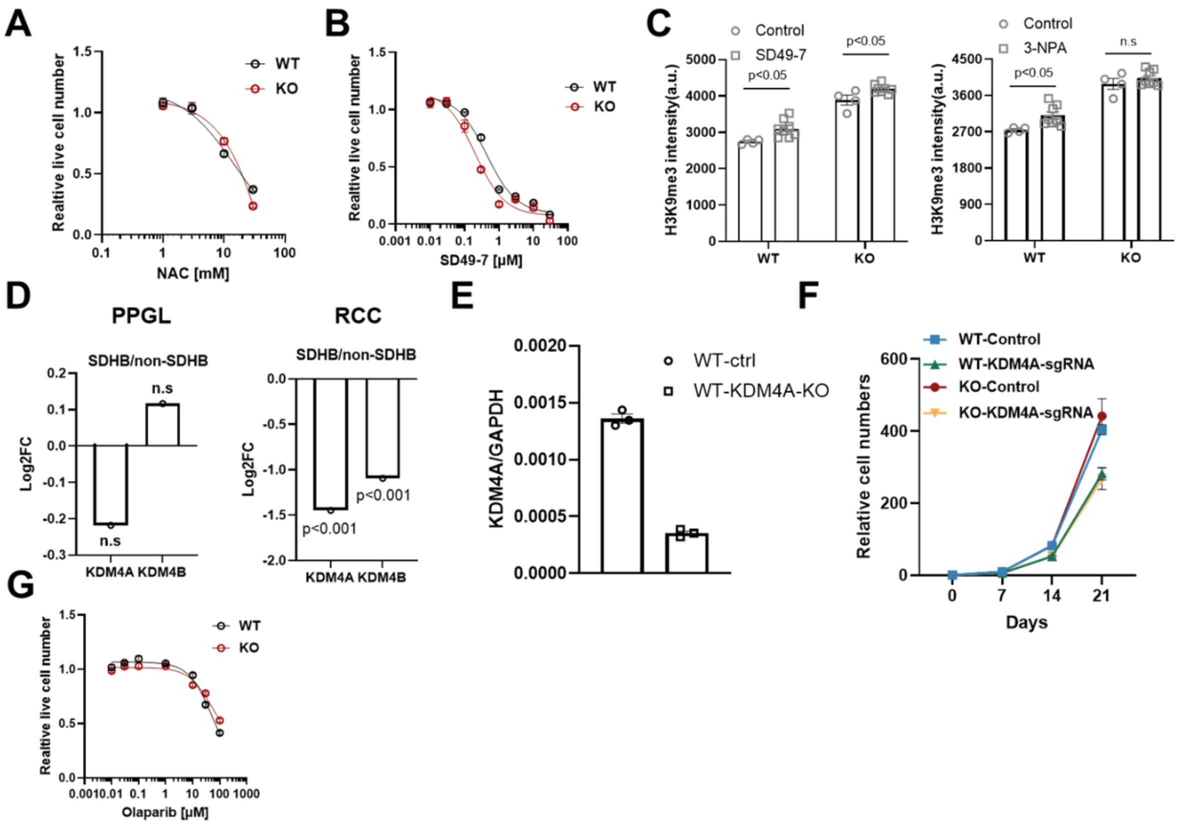


**Supplementary Fig. 5. Expression profile in *SDHB*-mutant tumors and the effect of disrupting KDM4A in UOK269 cells.** (A) and (B) Dose–response curves of NAC and SD49-7 in WT and KO cells. n=3 independent experiments. (C) Quantification of H3K9me3 immunofluorescence signals in WT and KO cells with or without SD49-7 (left) or 3-NPA (right) treatment. n=3 independent experiments. A linear mixed model analysis was applied in which treatment, and each independent experiment were used as the fixed effect and the random effect, respectively. (D) Comparison of KDM4A and KDM4B transcript levels between *SDHB*-mutant and non–*SDHB*-mutant PPGLs and renal cell carcinomas (RCCs) using TCGA datasets. (E) RT-PCR analysis of KDM4A in WT control and WT-KDM4A knockout cells. n=3 samples. (F) Growth curve of WT and KO cells with or without KDM4A expression. n=3 samples. (G) Dose–response curves of Olaparib-PARP inhibitor in WT and KO cells. n=3 independent experiments.
